# Genetic factors contributing to extensive variability of sex-specific hepatic gene expression in Diversity Outbred mice

**DOI:** 10.1101/2020.06.29.177311

**Authors:** Tisha Melia, David J. Waxman

## Abstract

Sex-specific transcription characterizes hundreds of genes in mouse liver, many implicated in sex-differential drug and lipid metabolism and disease susceptibility. While the regulation of liver sex differences by growth hormone-activated STAT5 is well established, little is known about autosomal genetic factors regulating the sex-specific liver transcriptome. Here we show, using genotyping and expression data from a large population of Diversity Outbred mice, that genetic factors work in tandem with growth hormone to control the individual variability of hundreds of sex-biased genes, including many lncRNA genes. Significant associations between single nucleotide polymorphisms and sex-specific gene expression were identified as expression quantitative trait loci (eQTLs), many of which showed strong sex-dependent associations. Remarkably, autosomal genetic modifiers of sex-specific genes were found to account for more than 200 instances of gain or loss of sex-specificity across eight Diversity Outbred mouse founder strains. Sex-biased STAT5 binding sites and open chromatin regions with strain-specific variants were significantly enriched at eQTL regions regulating correspondingly sex-specific genes, supporting the proposed functional regulatory nature of the eQTL regions identified. Binding of the male-biased, growth hormone-regulated repressor BCL6 was most highly enriched at *trans*-eQTL regions controlling female-specific genes. Co-regulated gene clusters defined by overlapping eQTLs included sets of highly correlated genes from different chromosomes, further supporting *trans*-eQTL action. These findings elucidate how an unexpectedly large number of autosomal factors work in tandem with growth hormone signaling pathways to regulate the individual variability associated with sex differences in liver metabolism and disease.

**Author summary:** Male-female differences in liver gene expression confer sex differences in many biological processes relevant to health and disease, including lipid and drug metabolism and liver disease susceptibility. While the role of hormonal factors, most notably growth hormone, in regulating hepatic sex differences is well established, little is known about how autosomal genetic factors impact sex differences on an individual basis. Here, we harness the power of mouse genetics provided by the Diversity Outbred mouse model to discover significant genome-wide associations between genetic variants and sex-specific liver gene expression. Remarkably, we found that autosomal expression quantitative trait loci with a strong sex-bias account for the loss or gain of sex-specific expression of more than 200 autosomal genes seen across eight founder mice strains. Genetic associations with sex-specific genes were enriched for sex-biased and growth hormone-dependent regulatory regions harboring strain-specific genetic variants. Co-regulated gene clusters identified by overlapping regulatory regions included highly correlated genes from different chromosomes. These findings reveal the extensive regulatory role played by autosomal genetic variants, working in tandem with growth hormone signaling pathways, in the transcriptional control of sex-biased genes, many of which have been implicated in sex differential outcomes in liver metabolism and disease susceptibility.

## Introduction

Sex differences in mammalian gene expression are not limited to reproductive tissues, but also occur somatic tissues [1], most notably the liver, as seen in mouse [2–4], rat [5, 6] and human [7, 8]. Sex-differential gene expression impacts hundreds of genes that control sex differences in liver function, including metabolism of drugs [9–11], other xenobiotics [12, 13] and fatty acids [14], as well as sex differences in liver disease [7, 15–18]. Sex differences in the liver transcriptome characterize protein-coding transcripts [6, 19], long noncoding RNAs (lncRNAs) [20, 21] and miRNAs [22–24]. Growth hormone (GH) is a key regulator of sex-biased gene expression [25, 26] and acts via its sex-dependent pattern of pituitary secretion: pulsatile in males and nearly continuous in females [27–29]. These sex differential plasma GH profiles, in turn, induce the sex-differential activation of the JAK2/STAT5 signaling pathway in hepatocytes. STAT5 and downstream GH-dependent transcription factors bind to individual sites in liver chromatin in a sex-biased manner, resulting in sex-differences in liver transcription [30–32]. Sex differences are also evident in the liver epigenome, including extensive sex differences in open chromatin regions (DNase I hypersensitive sites; DHS) [33, 34], chromatin marks [4, 35, 36], and DNA methylation [37–39] associated with the transcription of sex-specific genes.

Genetic modifiers can contribute to the regulation of sex-biased genes. In one early example, male-biased expression of *Cyp2d9* is abolished by a single nucleotide polymorphism (SNP) in the 5’ regulatory region [40]. In another example, the Regulator of Sex-Limitation (*Rsl*) gene represses certain sex-specific genes in a strain-specific manner [41, 42]. Genetic polymorphisms can impart individual differences in drug-metabolizing enzymes [43], many of which are encoded by sex-specific genes [11, 44]. Genome-wide association studies linking genetic variants and liver gene expression have identified many expression quantitative trait loci (eQTLs) for liver-expressed genes [45–49]. Little is known, however, about the extent to which such genetic factors impact individual variability in sex-biased gene expression. Understanding the source of variability in sex-biased gene expression in liver has far-reaching implications, as sex-biased genes can impact liver diseases with sex differences in susceptibility, severity and in some cases etiology [50], including non-alcoholic fatty liver disease (NAFLD) [51] and hepatocellular carcinoma [52, 53], and polygenic dyslipidemia and coronary artery disease [7, 54, 55].

Here, we investigate the impact of genetic regulation on sex-specific gene expression using Diversity Outbred (DO) mice, an outbred population derived from eight inbred mouse strains [56, 57]. DO mice have high natural allelic variance [58, 59] and diverse phenotypes [60, 61], which facilitates fine genetic mapping of liver-specific traits. Full genomic sequences are available for all eight DO founder strains [62], allowing genotypes in individual DO mice to be mapped back to specific founder strains. Using this model, we find unexpectedly high rates of gain and loss of sex-specific gene expression across DO founder mouse strains. We identify novel autosomal eQTLs targeting thousands of liver-expressed genes, including 1,500 eQTLs regulating liver-expressed lncRNA genes. Almost 1,000 of the eQTLs we identified target sex-specific genes, and many of these show sex-dependent genetic associations that explain the gain or loss of sex-specific gene expression in DO founder mice. Further, we link sex-specific binding of GH-regulated transcription factors and sex differences in chromatin accessibility to eQTL regions regulating sex-specific genes, and we identify co-regulated gene clusters defined by their overlapping eQTLs, a subset of which involve *trans* regulation. This elucidation of genetic factors controlling liver sex differences has important implications for our understanding of inter-individual variability in sex-biased liver function and disease.

## Results

### High variability of sex-specific gene expression across DO mouse founder strains

We identified 1,033 protein-coding genes that show sex-specific expression in mouse liver in one or more of nine mouse strains examined (418 male-biased genes and 615 female-biased genes). These strains include CD-1 mice, used in our earlier studies of sex-biased liver gene expression [34], and the eight founder strains used to establish the DO mouse population (A/J, C57BL/6J, 129S1/SvlmJ, NOD/ShiLtJ, NZO/HILtJ, CAST/EiJ, PWK/PhJ, WSB/EiJ) [56, 59]. Sex-biased gene expression was highly variable across the eight DO founder strains (Fig. 1A), with sex bias lost in at least one strain for 96% of the genes examined, and only 20 genes showing sex-biased expression in all 8 strains (see full dataset in Table S1; examples are shown in Fig. S1). This strong strain dependence also characterized the most highly sex-specific genes, with sex-biased expression lost in at least one strain for 58 of 74 genes with a sex-specific expression ratio > 4 at FDR < 0.05. Finally, analysis of the 220 sex-specific genes with the highest inter-strain variability revealed expression patterns highly specific to each strain, further confirming the striking variability of sex-specific gene expression in DO founder mouse livers (Fig. 1B).

**Fig. 1.**
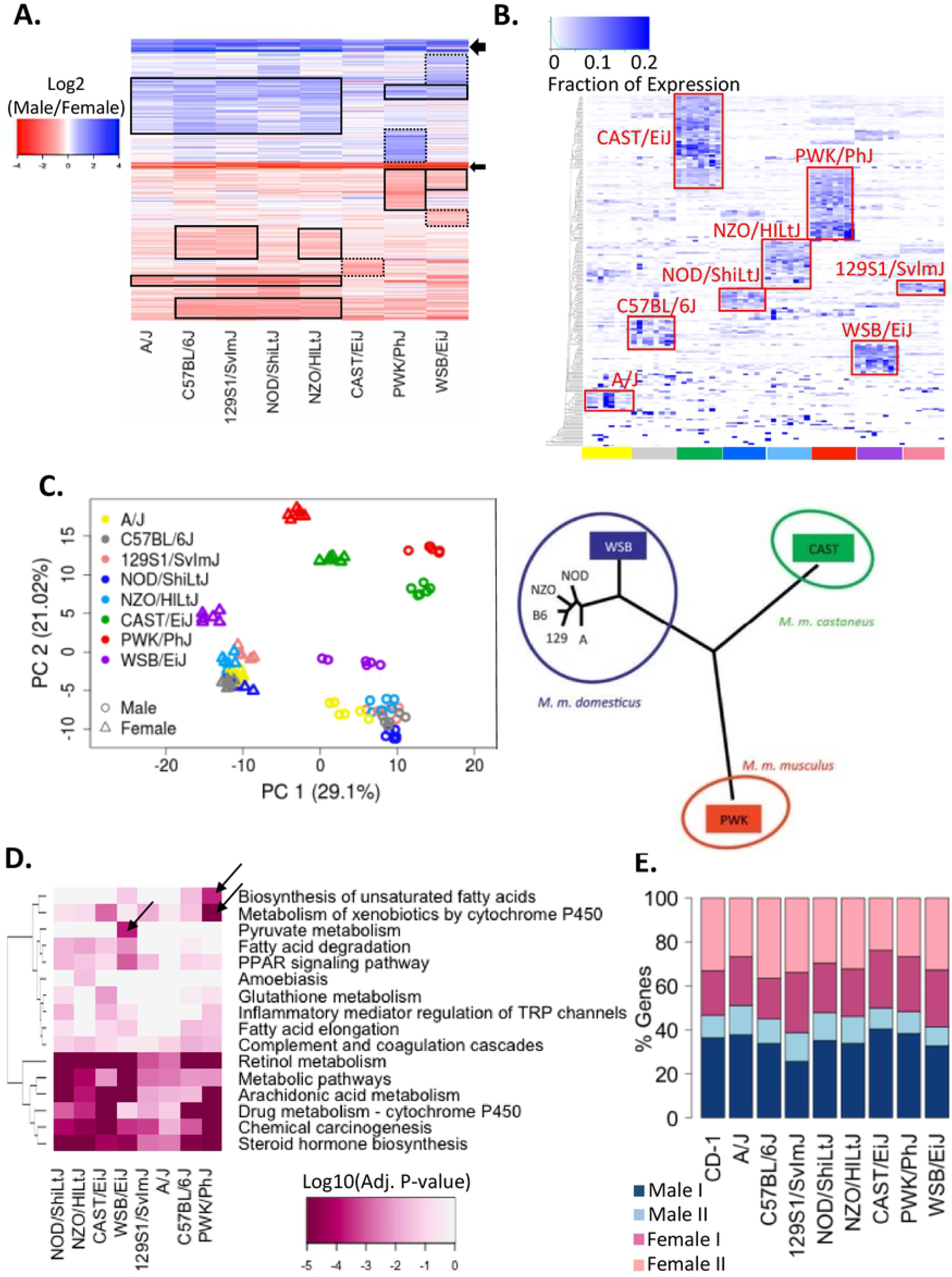
Sex-specificity across DO founder strains. **(A)** Heatmap of log2 male/female fold-change values across DO founder mouse livers for 471 protein-coding genes that show sex-biased expression (male/female |fold-change| > 1.5 at FDR < 0.05) in at least one DO founder strain based on microarray analysis. Solid black boxes, genes with strong sex-specificity in the indicated strains; dashed boxes, strong sex-specificity is limited to one strain. Arrows, strong sex-biased expression in all 8 strains. Sex-specific expression was lost in at least one of the 8 strains for 451 of these 471 genes. **(B)** Gene expression level for 220 sex-specific genes with the highest ratio of inter-strain variability to intra-strain variability (see Methods) across DO founder mouse livers. The map was clustered based on the summation of expression levels of each gene across the 8 indicated DO founder strain male livers, with the color intensity indicating the fraction of expression seen in each liver. Data shown are based on 8 male livers for each DO founder strain. **(C)** PCA of 96 microarray datasets (6 livers/sex/strain) based on the expression level of 471 sex-specific genes in each DO founder strain. Shown at the right is a phylogenetic tree [63] of the eight DO founder strains based on the SNP data for chr11. The three DO founder strains whose names are in colored boxes are the more distant wild-derived inbred strains, which matches the more distant patterns of sex-specific gene expression seen in the PCA plot. **(D)** KEGG pathways enriched (adjusted Benjamini P < 0.05) in the set of sex-specific protein-coding genes identified in each DO founder mouse strain. **(E)** Distribution of male-specific class 1 and class 2 genes, and of female-specific class 1 and class 2 genes across mouse strains. Each bar presents the distribution of sex-specific genes across four hypophysectomy response classes identified earlier [30].

The number of sex-specific genes identified in each founder strain ranged 2.4-fold, from 77 genes (strain A/J) to 183 genes (strain PWK/PhJ; Fig. S2A, Table S1). A similar pattern was seen when only the most highly sex-specific genes were considered (Fig. S2B). Principal component analysis (PCA) based on the expression levels of all sex-specific genes revealed that the largest variance (PC1) corresponds to the sex of each liver, as expected, while the second component (PC2) clustered the individual liver samples based on the evolutionary distance between strains (Fig. 1C). Thus, the six *Mus musculus domesticus* subspecies strains are closer to each other than to the other subspecies strains, i.e. *Mus musculus castaneus* (strain CAST/EiJ) and *Mus musculus musculus* (strain PWK/PhJ). Further, among the *Mus musculus domesticus* strains, the lone, wild-derived WSB/EiJ strain was separated from the other five strains, recapitulating the published phylogenetic tree shown in Fig. 1C [63].

We implemented DAVID analysis [64, 65] to determine whether the sex-specific genes identified in each strain showed differential enrichment for particular biological pathways. Sixteen KEGG pathways were enriched (adjusted P < 0.05) in at least one of the eight sets of sex-specific genes examined (Fig. 1D). Six pathways were significantly enriched in nearly all strains, including cytochrome P450-catalyzed drug and lipid metabolism, identifying this as a conserved sex-specific biological function, while ten pathways showed strain-specific enrichments (see y-axis dendrogram). Pyruvate metabolism was uniquely linked to sex-specific genes in WSB/EiJ livers, while biosynthesis of unsaturated fatty acids was most strongly enriched in PWK/PhJ livers.

### Impact on GH regulatory mechanisms

Our finding that some mouse strains contain comparatively few sex-specific genes, e.g., A/J and CAST/EiJ mice (Fig. S2), raises the possibility that these strains might be deficient in one of the four major classes of GH-regulated sex-specific genes found in liver [30, 66]. These classes of sex-specific genes are defined based on their responses to surgical removal of the pituitary gland by hypophysectomy, which ablates pituitary secretion of GH, as well as secretion of all other pituitary-dependent hormones. Class 1 male-specific and female-specific genes are activated in liver by the pituitary GH secretion profile in livers of the sex where they are more highly expressed, and consequently, following hypophysectomy, class 1 genes are down regulated in the sex where they show higher expression. In contrast, class 2 male-specific and female-specific genes are repressed in liver by the GH secretion profile of the sex where they are less highly expressed, and consequently, following hypophysectomy, class 2 genes are up regulated (de-repressed) in the sex where they show lower expression. We did not observe any major differences in the distribution of sex-specific genes across the four GH/hypophysectomy response classes in any of the DO founder strains, as compared to the distribution in CD-1 mouse liver (Fig. 1E). Thus, the major GH-dependent regulatory mechanisms for liver sex-biased gene expression captured by the hypophysectomy class 1/class 2 gene designations are apparently preserved across the DO founder strains, despite high variability in the expression seen for many individual sex-specific genes.

### eQTLs for liver-expressed genes

We analyzed published genotyping and expression datasets from 438 individual DO mouse livers [45, 67, 68] (Table S2) to discover eQTLs representing significant associations between SNPs/indels and the expression of either protein coding genes or multi-exonic liver-expressed lncRNA genes [20, 21]. This analysis was performed using the full set of 438 livers, and separately, all male livers (n=219) or all female livers (n=219) (Fig. S3A). We identified 10,325 significant autosomal eQTLs when considering all livers, 7,414 autosomal eQTLs when considering male livers only, and 8,233 autosomal eQTLs when considering female livers only, giving a total of 12,886 autosomal eQTLs (Table 1, Fig. 2A). The total number of eQTLs increased to 17,278 were found when chrX eQTLs were included (Table S3, Table S4A). Half of the autosomal eQTLs identified (6,344 out of 12,886) were not identified in earlier studies [45, 47–49, 68] and are novel, and include a large majority of the 1,524 eQTLs associated with liver-expressed lncRNA genes [20]. More autosomal eQTLs were identified in female than in male liver (Table 1), similar to findings in BXD mouse liver [48], suggesting genetic regulation is more common for the female liver transcriptome. In striking contrast, male DO liver eQTLs on chrX were 8.4 times more frequent than female DO liver eQTLs (3,932 vs. 468 eQTLs; Table S3A). This latter finding is consistent with the enrichment of sex-interacting eQTLs on human chrX [69], and with the observation that X-linked genetic variation in humans is significantly greater in males than in females [70]. Finally, 66% of autosomal eQTLs are within the same TAD [71] as the genes they are associated with, consistent with genetic regulation in *cis* (Table 1). In contrast, all 4,392 eQTLs on chrX were *trans*-acting (Table S4A).

**Fig. 2.**
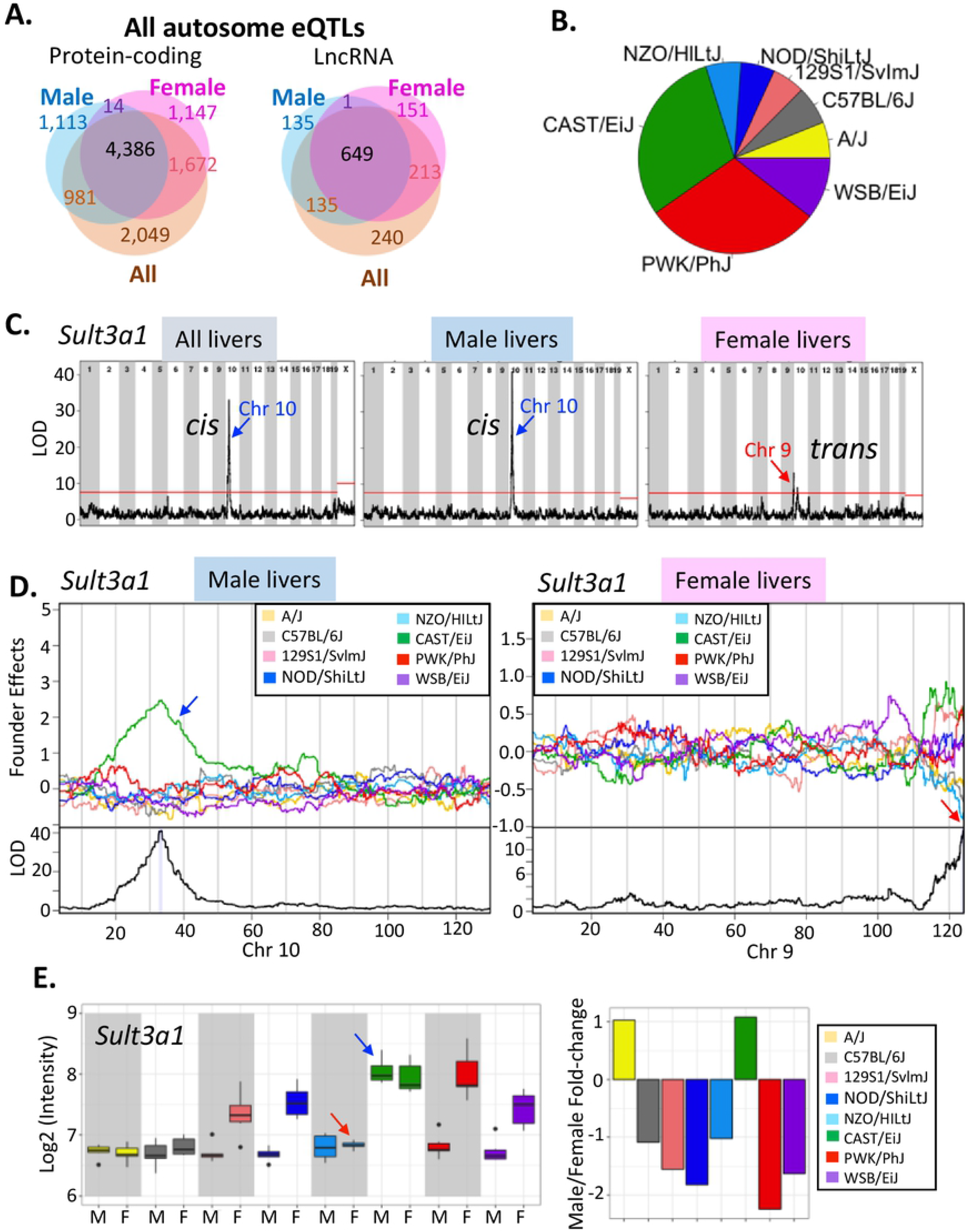
Autosomal eQTLs for liver-expressed genes. **(A)** Venn diagrams showing numbers of autosomal eQTLs discovered using all DO livers, or using male only or female only liver samples and that are associated with protein-coding genes (*left*) or lncRNA genes (*right*). Overall, 10,597 autosomal eQTLs are significant in either male liver or female liver (Table 1). **(B)**. Pie chart showing the realtive frequency at which each of the indicated DO founder strains has the largest regression coefficient (i.e., is the major regulating strain) for the combined set of 12,886 autosomal eQTLs. **(C)** Genome-wide association of *Sult3a1* in all, male only, and female only DO livers. The horizontal red line marks the P < 0.05 significance cutoff based on the permutation test. **(D)** Regression coefficients (*top* of each panel) and LOD scores (log10(p-value); *bottom* of each panel) across the chromosome that has a significant eQTL peak in male (*left*) or female (*right*) mouse liver, as marked at bottom. Shaded area in the LOD score plot indicates the 95% Bayesian credible interval for each eQTL. **(E)** Expression of *Sult3a1* in male and female mouse livers for each DO founder strain (*left*). Also shown are calculated male/female expression ratios (*right*). The middle hinge of each boxplot indicates the median, and the lower and upper hinges indicate the first and third quartiles. Whiskers mark 1.5 * IQR value. Red arrows indicate a decrease in female liver expression in NZO/HiLtJ mice and an increase in male liver expression in CAST/EiJ mice, both of which abolish the sex-specific expression seen in the respective mice strain.

**Table 1.** Number of autosomal eQTLs found when using all, male only or female only DO mouse livers. The number of genes associated with each set of eQTLs is shown in parenthesis. Last column: number of Combined eQTLs found in the same TAD as the genes they are associated with. More significant eQTLs were found in female than male DO mouse liver, similar to BXD mouse liver, where 1,638 significant eQTLs were found in male liver vs. 2,076 significant eQTLs in female liver (Gatti DM et al, 2010). See Table S4A for full listing of eQTLs.

|  | Set of genes | All Livers | Male Livers | Female Livers | Combined | Cis eQTLs |
| --- | --- | --- | --- | --- | --- | --- |
|  |  | number of eQTLs (number of genes) |  |  |  |  |
| All eQTLs | Protein-coding<br>(n = 18,543) | 9,088<br>(8,753 genes) | 6,494<br>(6,202 genes) | 7,219<br>(6,904 genes) | 11,362<br>(10,097 genes) | 7,492<br>(66 %) |
|  | LncRNA<br>(n = 2,016) | 1,237<br>(1,177 genes) | 920<br>(885 genes) | 1,014<br>(957 genes) | 1,524<br>(1,318 genes) | 1,014<br>(67%) |
|  | **Total eQTLs:** | **10,325** | **7,414** | **8,233** | **12,886** | **8,506** |
| eQTLs<br>associated<br>with 1,196<br>sex-specific<br>genes | Protein-coding<br>(n = 1,033) | 708<br>(682 genes) | 555<br>(523 genes) | 602<br>(581 genes) | 830<br>(743 genes) | 631<br>(76%) |
|  | LncRNA<br>(n = 163) | 131<br>(121 genes) | 99<br>(95 genes) | 105<br>(97 genes) | 157<br>(130 genes) | 108<br>(69%) |
|  | **Total eQTLs:** | **839** | **654** | **707** | **987** | **739** |

For each eQTL, we identified the founder strain whose SNP/indel is most likely responsible for the genetic association captured by the eQTL (the ‘regulating strain’). This was indicated by the founder strain whose regression coefficient at the SNP marker where the strongest association occurs (i.e. the SNP marker with the highest LOD score) had the largest absolute value (Table S4A). 60% of all autosomal eQTLs discovered using all DO livers (6,158 of 10,325 eQTLs) were associated with genetic variants in either CAST/EiJ or PWK/PhJ mice (Fig. 2B). Those two strains are the most evolutionarily divergent (Fig. 1C) and have the largest number of SNPs and indels [59]. The WSB/EiJ strain follows (10% all of autosomal eQTLs), followed by the other five founder strains (~6% of autosomal eQTLs each), highlighting how liver gene expression patterns diverged as genetic variants accumulated during evolution. A majority (54-67%) of all autosomal eQTLs were associated with up regulation of gene expression in the regulating strain (Fig. S3B), and 36% (3,706 of 10,325 autosomal eQTLs) were associated with changes in expression in two or more regulating strains (Table S4A, column AC).

### Genetic regulation of sex-specific genes

eQTLs regulate a large fraction of sex-specific genes, with 873 of 1,196 sex-specific genes (73%) being associated with a total of 987 autosomal liver eQTLs (Table 1, Fig. 3A). 143 of these eQTLs were only significant when using male livers (66 eQTLs) or only when using female livers (77 eQTLs) (Table S4A). Further, many of the eQTLs discovered using both male and female DO livers showed a stronger genetic association in one sex (Table S4B, Fig. S4). eQTLs that are stronger in male DO livers were biased for association with strongly male-specific genes, and those that are stronger in female liver were more closely associated with strongly female-specific genes, as indicated by the distributions of their sex-differential LOD scores (Fig. 3B). eQTLs within the top 10% of LOD score differences between male and female livers (top 1,060 of all 10,597 such autosomal eQTLs; Table S4A) were enriched 5.1-fold for association with genes of matched-sex-specificity, as compared to genes with the opposite sex-specificity (P-value < 6.5e-07, Fisher’s Exact test). Multiple eQTLs were identified for 199 sex-specific genes (Table S4C), indicating these genes are subject to multiple, distinct forms of regulation. Finally, 240 sex-specific genes were not associated with any eQTLs (Table S4D). These expression of these 240 genes across DO livers was less variable than that of sex-specific genes with eQTLs, as was seen in both male DO (P < 0.01, Student’s t-test) and female DO livers (P < 0.007, Student’s t-test); average expression levels were similar for both gene sets (P> 0.28, Student’s t-test).

**Fig. 3.**
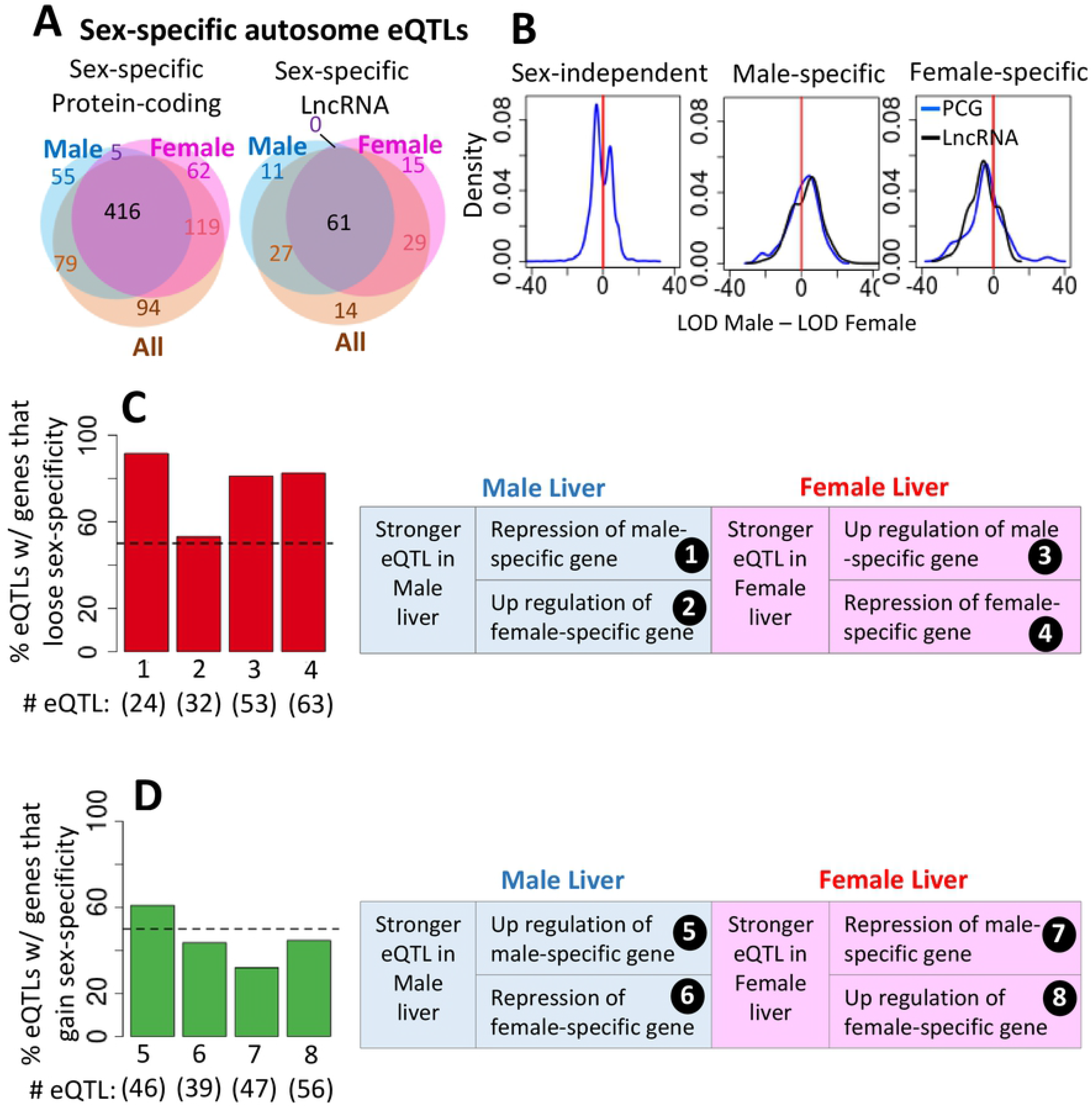
Autosomal eQTLs associated with sex-specific genes. **(A)** Venn diagrams showing overlaps for numbers of eQTLs discovered in all, male only or female only DO liver samples that are associated with sex-specific protein-coding or sex-specific, multi-exonic intergenic lncRNA genes. **(B)** Distribution of LOD score differences (log10 values) in male vs. female liver for autosomal eQTLs that are associated with genes that are sex-independent (*left*), strongly male-specific (male/female |fold-change| > 4; *middle*) or strongly female-specific (male/female |fold-change| > 4; *left*). **(C)** (*Left*) Distribution of 172 eQTLs in categories #1-4 (described at the right), which are associated with a loss of sex-specific gene expression in the regulating strains. (*Right*) Four scenarios, whereby an eQTL deceases the sex-specfiicity of gene expression. **(D)** (*Left*) Distribution of 188 eQTLs in categories #5-8 (described at the right), which are associated with a gain of sex-specific gene expression in the regulating strains. (*Right*) Four scenarios, whereby an eQTL increases the sex-specificity of gene expression.

Next, we investigated whether eQTLs with a sex-bias in their association can explain the variability in sex-specific gene expression seen in DO founder strains. We examined 360 autosomal eQTLs for sex-specific genes that are significant in either male only or female only DO liver samples (or both) and are associated with either an increase or a decrease in sex-specificity in the regulating strain (Table S5, Table S6). These associations encompass 322 (68%) of the 471 sex-specific protein-coding genes for which expression data is available for all eight founder strains in both sexes. We identified four circumstances whereby an eQTL decreases the sex-specificity of expression in the regulating strain (Fig. 3C; 172 eQTLs in categories # 1-4), and four other circumstances whereby an eQTL increases sex-specificity in the regulating strain (Fig. 3D; 188 eQTLs in categories # 5-8) (Table S6, Fig. S5), where a decrease or an increase in expression is inferred from the regression coefficient in the regulating strain (Table S5). One example is the category # 2 eQTL on chr10 regulating the female-specific *Sult3a1* gene, which is highly significant in male DO livers but was not found in female livers (Fig. 2C). This *cis* eQTL is associated with loss of sex-specificity due to the elevated expression of *Sult3a1* in male livers of CAST/EiJ mice, the regulating strain (Fig. 2D, *left*; Fig. 2E*; blue arrows*). *Sult3a1* is also associated with a distinct, category # 4 eQTL on chr9 that was uniquely identified in female livers (Fig. 2C). This *trans* eQTL abolishes sex-specificity due to the decreased expression of *Sult3a1* in female livers of NZO/HIltJ mice, its regulating strain (Fig. 2D, *right*; Fig. 2E; *red arrows*). Thus, both eQTLs abolish the sex difference in *Sult3a1* expression in their respective regulating founder strains (Fig. 2E, *arrows*), but by different mechanisms.

Overall, 219 of the 360 eQTLs (61%), impacting 205 sex-biased genes, were associated with either an outright loss or gain of sex-specificity in the regulating strain (Table S6, column D). For 134 of the 172 (78%) eQTLs in categories # 1-4, sex-specific gene expression is lost in the regulating strain but it is retained in at least one other strain. Sex-specific expression is lost in the regulating strain for 81-92% of eQTLs in categories # 1, 3 and 4, but for only 53% of category # 2 eQTLs (Fig. 3C). This indicates that while all category # 2 eQTLs decrease sex-specificity in the regulating strain, almost half are not strong enough to cause a loss of sex-specificity. Of the 188 eQTLs in categories # 5-8, only 47% (85 eQTLs) showed a gain of sex-specificity in the regulating strain (Fig. 3D). This suggests there is another layer of gene regulation for many of those genes. These findings highlight the extensive impact and multiple mechanisms through which genetic factors regulate the variability of sex-specific gene expression in DO mice.

### Sex-biased regulatory elements within eQTL regions

Genomic regions encompassed by an eQTL may impact sex-specific gene expression in the regulating founder strain through strain-specific genetic variants at sex-specific regulatory regions, e.g., enhancers. Such regulatory regions include DHS that are sex-biased in mouse liver, i.e., regions where chromatin is significantly more open (more accessible) in one sex, and that harbor sex-biased binding sites for key GH-regulated liver transcription factors [31, 32, 35]. Strain-specific genetic variants at such sites may alter or abolish GH-regulated transcription factor binding activity, and perhaps may alter chromatin accessibility and thereby dysregulate the expression of sex-biased genes. To investigate this hypothesis, we analyzed sex-biased DHS, and separately, sex-biased transcription factor binding sites that contain SNPs/indels specific to the identified regulating strain within each eQTL region (Table S4A, last columns). Sex-biased DHS with strain-specific variants were enriched at eQTL regions associated with genes with matching sex-specificity, as compared to eQTLs associated with genes showing the opposite sex-bias or genes with no sex-specificity (Fig. 4A; *first two panels*; Fig. S6). The majority of DHS found in any eQTL region, both for sex-biased and sex-independent genes, are sex-independent DHS; these DHS showed a significant enrichment at eQTLs for sex-independent genes, as compared to sex-specific genes (Fig. 4A; *third panel*). A similar pattern of enrichment was found for sex-biased binding sites for the GH-regulated transcription factor STAT5 (Fig. 4B). In contrast, male-biased binding sites for the GH-regulated transcriptional repressor BCL6, which preferentially represses female-specific genes in male liver [32, 72], showed significant enrichment at *cis* eQTL regions for female-specific genes, when compared to eQTL regions for male-specific or sex-independent genes (Fig. 4C, *first panel*). Sex-independent BCL6 binding sites were significantly depleted at eQTL regions associated with female-specific genes (Fig. 4C, *third panel*). Male-biased BCL6 binding sites showed an even greater enrichment at *trans* eQTL regions (Fig. 4D). These enrichments of sex-biased DHS and transcription factor binding sites with strain-specific SNPs/indels provide strong support for the functional relevance of these genomic regions to the eQTL-based regulation.

**Fig. 4.**
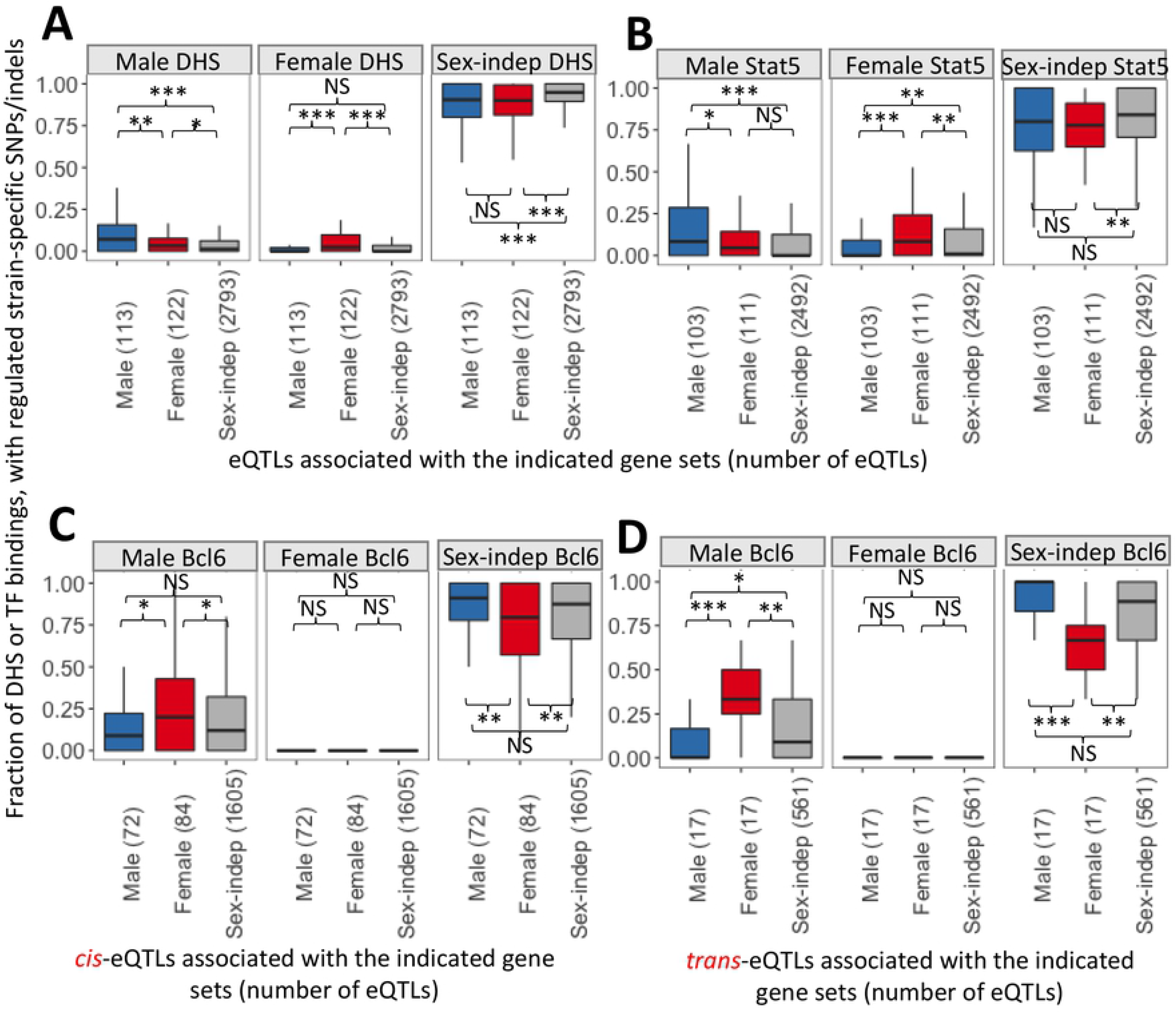
Enrichment at different sets of eQTL regions of sex-biased DHS and transcription factor binding sites that contain regulating strain-specific SNPs/indels. Y-axis, fraction of all DHS (A), or all binding sites for STAT5 (B) or BCL6 (C, for *cis*-eQTLs; and D, for *trans*-eQTLs) within the eQTL region that are male-specific, female-specific or sex-independent and that contain (within the DHS or within the binding site) SNPs/indels specific to the regulating strain. This analysis was carried out for the sets of eQTLs for genes expressed in a male-specific, female-specific, or sex-independent manner (sets of eQTLs specified along the x-axis), with the number of eQTLs analyzed in each boxplot indicated in parenthesis. Data shown in A and B are for *cis* plus *trans* eQTLs, whereas *cis* (C) and *trans* (D) eQTLs were analyzed separately in C and D. The number of DHS and STAT5 or BCL6 binding sites in each eQTL is indicated in Table S4. For example, each of the three red boxplots in panel D shows the distribution of 17 values, where each value is the fraction of BCL6 binding sites that are male-biased, female-biased, or sex-indepenent (as indicated at top) in a set of 17 *trans* eQTL regions that are associated with female-specific genes; only BCL6 binding sites that contain SNPs/indels that are specific for the regulated strain are considered. For each factor, the distributions were compared across eQTL sets using the Wilcoxon test (***, p<0.001, **, p<0.01 and *, p<0.05), as indicated in each subpanel. The middle hinge of each boxplot indicates the median value, while the lower and upper hinges correspond to the first and third quartiles. Whiskers marks 1.5 * IQR value.

### Inference of gene co-regulation based on eQTLs

We hypothesized that genes with overlapping eQTLs are likely to be co-regulated, presumably via shared regulatory regions. Overlapping eQTLs with a common regulating strain were used to define co-regulated gene clusters. We identified 1,521 such co-regulated gene clusters, comprising 4,105 genes, based on eQTLs discovered in all DO liver samples (Table S7A). The vast majority of the co-regulated gene clusters are small in size; 95% of all clusters include five or fewer genes each, although some clusters contain as many as 25 genes (Table S7D). 28 of the gene clusters were significantly enriched for sex-specific genes (Table S7A). The absence of many large co-regulated gene clusters indicates that strain differences are primarily determined by many individual loci, each regulating a small number of genes, rather than by a small number of variants in master regulatory regions. The gene clusters we identified show evidence of co-regulation, with many genes within clusters showing high pairwise gene expression correlation (see examples in Fig. 5). Half (762) of the co-regulated clusters contain sub-clusters, where the identified eQTL region was associated with up regulation of some genes and down regulation of other genes in the cluster. One example is a 21-gene cluster on chr2, which is predicted to occur in the PWK/PhJ strain (Fig. 5A; Table S7A, Cluster100_all). Pairwise gene expression correlation analysis separated these 21 eQTLs into two well-defined sub-clusters. Eleven overlapping eQTLs in the upper sub-cluster have positive regression coefficients (i.e. increased expression of the 11 genes; Fig. S7) and are all located on the same chromosome (chr2) as their associated genes, whereas 10 overlapping eQTLs in the lower sub-cluster have negative regression coefficients (i.e., decreased expression of the 10 genes; Fig. S7 and Fig. S8) and are each located on a different chromosome than the genes they are associated with. The pairwise gene correlations are less tight in the negatively correlated sub-cluster (e.g. *Zbtb7c*), suggesting the eQTL association may represent indirect regulation or partial regulation involving another factor. Notably, in the case of the eQTL involving *Zbtb7c*, an independent, stronger genetic association was identified in *cis* (chr18, Fig. S8). The co-regulated eQTL cluster in Fig. 5A shows that genes from different chromosomes may be grouped together (Fig. S7), highlighting the ability of this approach to identify complex *trans* regulation. Overall, 159 (10.4%) of the co-regulated clusters grouped genes from multiple chromosomes. Further, 344 (23%) of the co-regulated clusters contain both protein-coding and lncRNA genes, generating testable hypothesis for possible functions of the co-clustered lncRNAs. We also found examples of co-regulated clusters containing genes within the same chromosome, indicating local regulatory regions (Fig. 5B).

**Fig. 5.**
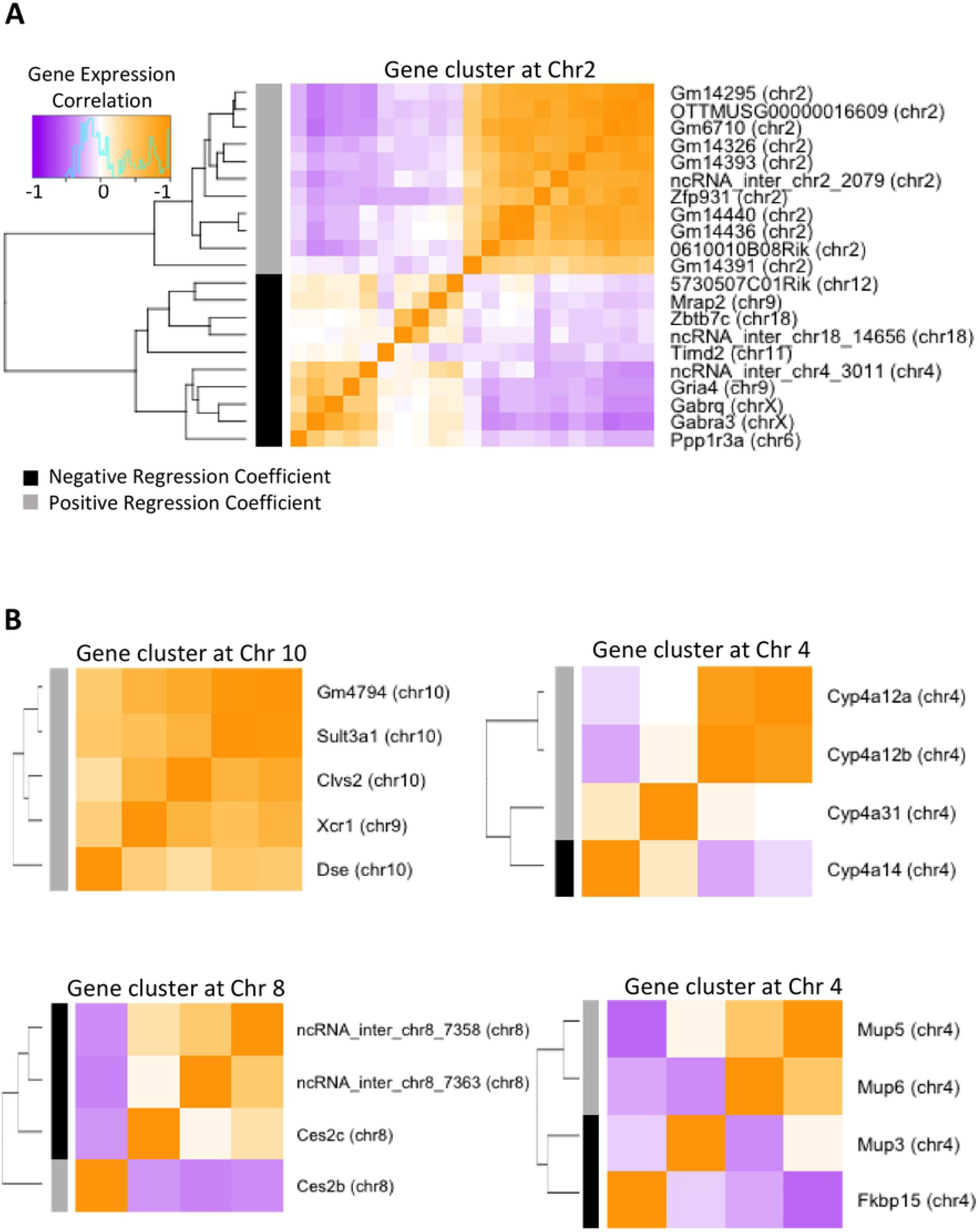
Co-regulated gene clusters based on overlapping eQTL regions discovered in all liver samples. **(A)** A co-regulated cluster consisting of twenty-one overlapping eQTLs with the PWK/PhJ as the regulating strain. **(B)** Four co-regulated clusters containing sex-specific genes. The heatmaps show Pearson correlations of gene expression.

Co-regulated gene clusters based on overlapping eQTLs were also discovered using male only DO livers (Table S7B), and separately, female only DO livers (Table S7C), and clusters with a sex-dependent genetic association were identified. Some of these clusters showed a switch in gene members between clusters identified in male liver as compared to female liver samples. One example involves three female-specific genes from the sulfotransferase gene family: *Sult2a1*, *Sult2a3* and *Sult2a5* (Fig. 6A). In PWK/PhJ male mouse liver, but not in PWK/PhJ female mouse liver, *Sult2a1* and *Sult2a5*, but not *Sult2a3,* are predicted to be co-regulated, as their eQTLs overlap and are associated with high expression of both genes (Fig. 6B, male samples, *top* and *bottom*). In contrast, in CAST/EiJ female liver, *Sult2a1* shows co-regulation with *Sult2a3*, and the high expression of both genes is associated with overlapping eQTL regions (Fig. 6B, female samples; *top* and *middle*). Even though all three eQTL regions overlap, their eQTL peaks do not overlap, indicating a need for higher resolution genetic mapping to pinpoint the precise regulatory regions and tease apart these two distinct regulations. *Sult2a3* and *Sult2a5* showed no significant eQTLs in male and female liver samples, respectively (Fig. S9). The pairwise gene correlations between these three genes are consistent with these patterns, as *Sult2a1* and *Sult2a3* show higher correlation in female liver samples, as compared to male liver samples (Pearson correlation 0.7 vs. 0.3), and *Sult2a1* and *Sult2a5* show higher correlation in the male samples, as compared to female samples (Pearson correlation 0.8 vs 0.6) (Fig. 6A). The relatively strong correlation in female livers between *Sult2a1* and *Sult2a5*, which was not seen in the CAST/EiJ co-regulated cluster, indicates that there is, nevertheless, an overall correlation in their expression across female DO mouse livers, once the complexity of other DO founder strains is factored in.

**Fig. 6.**
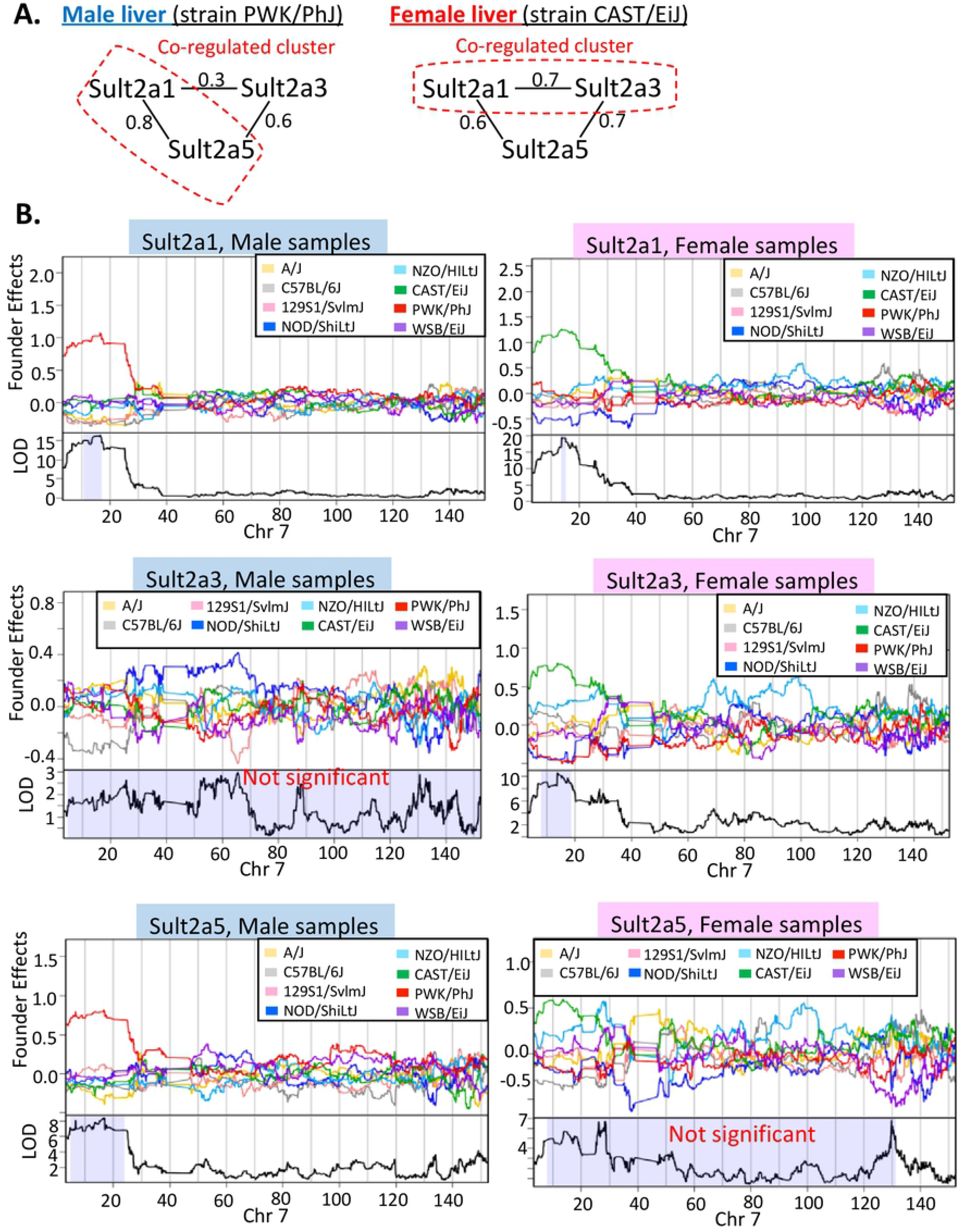
Co-regulated gene clusters that show different gene members in male vs. female liver. (**A)** Pearson gene expression correlation of *Sult2a1*, *Sult2a3* and *Sult2a5* in male liver (*left*) and female liver (*right*). **(B)** Regression coefficients (top figure in each panel) and LOD scores (bottom figure in each panel) across chr7 for *Sult2a1* in male liver (*top left*) and female liver (*top right*), for *Sult2a3* in male liver (*middle left*) and female liver (*middle right*) and for *Sult2a5* in male liver (*bottom left*) and female liver (*bottom right*). Shaded area in the LOD score figures indicate 95% Bayesian credible interval for each eQTL.

## Discussion

Sex-specific gene expression is widespread in mammalian liver, where it impacts male-female differences in lipid and drug metabolism as well as disease susceptibility. While the role of hormonal factors, most notably GH, in regulating liver sex differences is well established, little is known about the effects of genetic factors on the observed differences between the sexes. Here, we used the DO mouse model, a genetically diverse population derived from eight inbred mouse strains [56], to dissect the genetic component of sex-biased transcriptional regulation. We discovered that an unexpectedly large number of distinct genetic factors work in tandem with the hormonal environment to regulate the transcription of sex-biased genes in the liver. DO mice showed high variability in sex-specific gene expression that reflects the high variability of expression seen in the eight founder strains, with 96% of sex-specific genes deficient in sex-biased expression in one or more strains. We identified 987 associations between autosomal genetic variants and expression of sex-biased genes (eQTLs for sex-biased genes), including 157 eQTLs associated with 130 sex-biased multi-exonic lncRNA genes [20]. Many of these eQTLs showed large differences in genetic association (LOD score) between male and female liver populations and were closely linked to the loss or gain of sex-biased gene expression in the predicted regulating founder mouse strain. Genomic regions that contain genetic variants specific to the regulating founder strain and show sex differences in chromatin accessibility, or in the binding of the GH-regulated transcription factor STAT5, were significantly enriched for eQTLs associated with genes with matching sex-specificity, supporting the proposed functional, regulatory nature of the eQTL regions we identified. Gene clusters defined by overlapping eQTLs identified sets of co-regulated genes, in some cases from different chromosomes, consistent with the *trans*-acting nature of a subset of the eQTLs described here.

### Variability of sex-specific genes across DO founder mice

We found exceptional high variability of sex-specific gene expression across DO founder mouse strains, with some strains expressing many more sex-specific genes than others, and with sex-specific expression absent in at least one founder strain for nearly all sex-specific genes. This suggests that, for the vast majority of sex-specific genes, conservation of the sex-specificity of a given pathway or function, such as cytochrome P450 (CYP)-mediated lipid and drug metabolism, may be more important than conservation of the sex specificity of individual genes. For example, although sex specificity was preserved across all eight DO founder strains for only 4 of 28 sex-specific *Cyp* genes (male-specific *Cyp2d9* and *Cyp4a12a*; female-specific *Cyp2a4* and *Cyp17a1*), six of the strains expressed at least 15 sex-specific *Cyps* each, and the other two strains expressed 9-11 sex-specific *Cyp* genes each (Table S1). However, strain-specific enrichment of certain KEGG pathways was also seen, e.g., pyruvate metabolism in WSB/EiJ mice and biosynthesis of unsaturated fatty acids in PWK/PhJ mice. Of the 20 genes that showed consistent sex-specific expression across all eight strains, eight genes were active in steroid and drug/foreign chemical metabolism (*Cyp*, *Gst*, *Fmo, Aox,* and *Hsd* family members), three were enzymes of fatty acid biosynthesis and metabolism (*Elovl3*, *Acot3*, *Cyp4a12a*), two were complement genes (*C8b*, *C9*), and two were involved in liver fibrosis (*Prom1*, *Rtn4;* [73, 74]). The conserved sex specificity of these 20 genes suggests they carry out essential, non-redundant sex-specific functions. The strain-dependent expression seen for the much larger number of other sex-specific genes is consistent with the diverse hepatic phenotypes, including differences in sex-biased liver disease susceptibility, seen in various mouse strains and in outbred individuals [56, 75].

Genetic variants accounted for a large fraction of the variability of sex-biased gene expression seen in DO mouse liver. Thus, 68% of sex-specific genes examined (i.e., 322 of 471 sex-specific genes for which we have DO founder expression data) were regulated by a set of 360 eQTLs predicted to either decrease or increase sex-specificity in the regulating founder strain. The effects of these eQTLs act by a variety of mechanisms, e.g., by suppression of a male-specific gene or activation of a female-specific gene in male liver, which both decrease sex specificity; and by increasing activation of a male-specific gene or increasing suppression of a female-specific gene in male liver, both of which increase sex specificity. Remarkably, the genetic modifiers of sex-specific genes that we identified explain more than 200 instances of gain or loss of sex-specificity across the eight DO mouse founder strains, highlighting the widespread impact of genetic regulatory factors on the variability of sex-specific gene expression.

DO founder mice exhibit diverse liver phenotypes, including differences in susceptibility to NAFLD [76], liver fibrosis [77] and hepatoxicity [78, 79], as well as differences in liver inflammatory responses [80], whose molecular mechanisms, involve, at least in part, sex-biased genes [17, 18, 81–84]. Notably, the sex-dependent eQTLs identified here include *cis*-acting eQTLs for several sex-specific genes implicated in NAFLD, such as *Cyp7b1*, a male-specific P450 that inactivates estrogenic and hepatotoxic sterols and is a key driver of NAFLD [85–90], *Gstp1*, a male-biased oxidative stress-protective glutathione transferase with high risk allelic variants for NAFLD in humans [91–93], and *Nox4*, a male-biased NADPH oxidase that increases hepatocyte oxidative stress, apoptosis and liver fibrosis [94–96]. We also identified both *cis* and *trans* sex-dependent eQTLs for *Sult3a1*, a female-specific hepatic sulfotransferase that protects from benzene genotoxicity [97]. The fact that a majority of genetic risk factors for complex diseases, including sex-biased NAFLD and insulin resistance [90, 98–100], are in non-coding regions, primarily at active DHS [101–104], supports the proposal that the underlying SNPs/indels for many of the sex-dependent eQTLs that we identified are in DHS and other such regulatory regions.

### Relationship between genetic variants and GH regulatory mechanisms

Many of the eQTLs for sex-specific genes described here were substantially stronger in one sex than the other, indicating the underlying SNPs/indels act in a sex-dependent manner. The underlying mechanisms for these effects likely involve transcriptional and epigenetic regulatory events controlled by GH [4, 34, 35], the key regulator of sex-specific gene expression in the liver. Global mechanisms of sex-specific hormonal regulation are apparently maintained in the DO founder strains, as indicated by their very similar distributions of sex-specific genes across four distinct sex-specific gene classes, defined by their responses to the loss of GH signaling following hypophysectomy [30, 66]. Moreover, sex-biased open chromatin regions (DHS) and sex-biased, GH-regulated STAT5 binding sites containing strain-specific variants were significantly enriched at eQTL regions regulating correspondingly sex-specific genes, while binding of the male-biased, GH-regulated repressor BCL6 [32, 72] was most highly enriched at *trans*-eQTL regions controlling female-specific genes. These findings support the proposal that SNPs/indels at these GH-dependent regulatory regions cause either a loss or a gain in sex-biased transcription factor binding, and thereby alter GH-regulated gene expression in a sex-dependent manner. In one model (Fig. 7A), a strain-specific SNP/indel at one of the sex-biased regulatory elements associated with an eQTL with a strong repressive effect on *Cyp2b9* expression in female 129S1/SvImJ liver (Fig. S5D) interferes with the binding of factors required for *Cyp2b9* transcription. In another model (Fig. 7B), an activating SNP/indel at one of the sex-biased regulatory regions is proposed to explain the de-repression of *Sult3a1* in CAST/EiJ male liver (Fig. 2C). Further studies, including mutation of individual strain-specific SNPs/indels found at each of the regulatory regions in these eQTLs will be required to fully evaluate these proposed mechanisms.

**Fig. 7.**
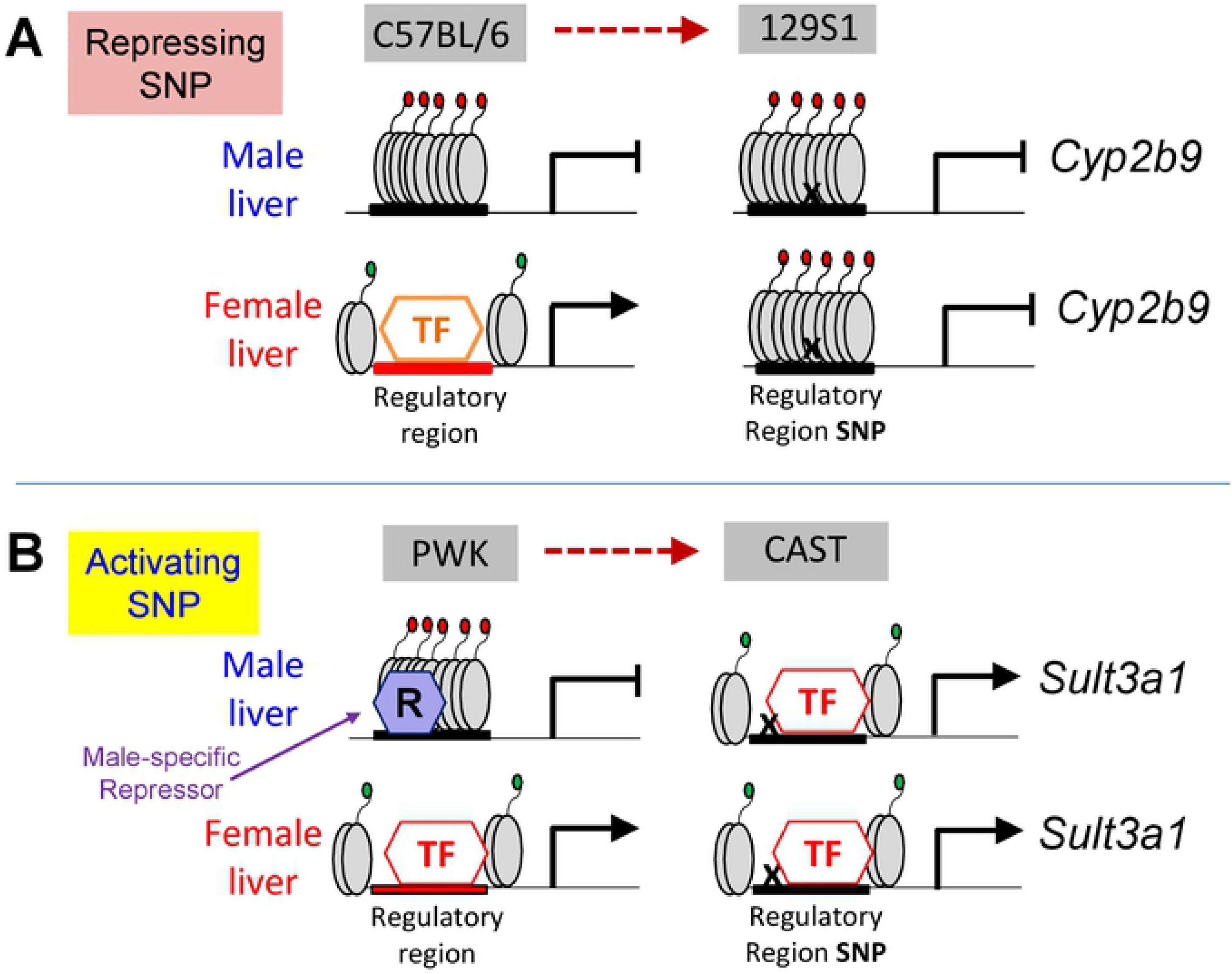
Models for how eQTLs may act in a sex-dependent manner to regulate sex-specific genes. **(A)** Repressing SNP at *Cyp2b9* regulatory element. A *cis*-acting eQTL on chr7 with a strong genetic association in female but not male mouse liver represses expression of *Cyp2b9* in female 129S1/SvImJ liver (Fig. S5D; category #4 eQTL, Fig. 3C). This eQTL is proposed to involve a SNP (‘X’) that abolishes binding of a female-specific transcriptional activator (TF) to the regulatory element, and may lead to loss of female-biased chromatin accessibility and the loss of the sex-specific expression seen in 129S1/SvImJ liver. The *Cyp2b9* eQTL region includes two female-biased DHS, three female-biased STAT5 binding sites, and five female-biased HNF6 binding sites, all of which overlap with 129S1/SvImJ-specific SNPs/indels (Table S4, Fig. S10). **(B)** Activating SNP at *Sult3a1* regulatory element. A *cis*-acting eQTL on chr10 with a much stronger genetic association in male than female mouse liver enables elevated expression of *Sult3a1* in male CAST/EiJ liver (Fig. 2C-2D; category #2 eQTL, Fig. 3C). This eQTL is proposed to involve a SNP (‘X’) that abolishes binding of a male-specific repressor protein (R) to the regulatory element shown, but does not interfere with binding of the transcriptional activator (TF), thereby causing loss of the sex-specific expression of *Sult3a1* in CAST/EiJ liver (Fig. 2E). The chr10 *Sult3a1* eQTL region includes two female-biased DHS, one female-biased STAT5 binding site and seven female-biased HNF6 binding sites [31], all of which overlap with 129S1/SvImJ-specific SNPs/indels (Table S4).

### Properties of the genetic regulation of sex-biased genes

Our findings give insight into several important features that govern how genetic modifiers impact the expression of sex-specific genes in the liver. We found major differences in the sex-specific liver transcriptome between mouse strains, determined by an unexpectedly large number of individual genetic loci, each regulating small numbers of genes in *cis* and/or in *trans*, and without affecting the overall patterns of GH regulation that dictate sex-specific gene transcription in the liver. Thus, the co-regulated gene clusters we identified from overlapping eQTLs in the same regulating strain are mostly small in size, containing five or fewer genes. However, this finding does not preclude the possibility that some of the clustered genes are themselves regulators of larger numbers of downstream genes, but whose effects are not sufficiently strong to be discovered as eQTLs in our analyses. Genetic modifiers of sex-specific gene expression may also operate via regulatory proteins that are not themselves expressed in a sex-dependent manner. One such example is *Rsl1*, which codes for a Krüppel-associated box (KRAB) zinc finger repressor whose expression varies across mouse strains [105, 106] and for which we found both *cis* and *trans* eQTLs (Table S4). *Rsl1* directly represses *Slp* and a few other male-specific genes in mouse liver, but can also regulate other sex-specific genes by indirect activation or repression, which enables genetic variants of *Rsl* to have a more widespread impact on sex-specific gene expression patterns [41, 42]. Another feature of genetic regulation seen here is the clustering of protein coding genes with co-regulated lncRNA genes, which may contribute to sex-specific gene regulation [20] through epigenetic or other regulatory mechanisms [81, 107, 108]. Finally, our finding of large numbers of sex-biased autosomal genetic associations in mouse liver differs from results in whole blood based on clinical samples, where a common autosomal genetic architecture for gene expression was found across the sexes [109].

## Materials and methods

### DO mouse liver datasets and analyses

Liver RNA sequencing expression data was downloaded from GEO (https://www.ncbi.nlm.nih.gov/geo/) for 438 individual DO mice using accession numbers GSE45684 [67, 68] and GSE72759 [45]. The datasets used for our analyses were derived from 219 male and 219 female DO mice, 26 weeks of age, all with SNP array data available. Of these, 112 males and 110 females were fed a standard chow diet, and 107 males and 109 females were fed a high fat diet. Genotyping data based on SNP arrays were downloaded from the DO Database (https://www.jax.org/research-and-faculty/genetic-diversity-initiative/tools-data/diversity-outbred-database#) for 264 DO samples genotyped at 7,854 SNPs using the Mouse Universal Genotyping Array (MUGA) [110] and for 174 DO samples genotyped using a higher density array, MegaMUGA [111], encompassing 77,725 SNP probes.

#### Haplotype reconstruction

Quantile normalization, implemented in DOQTL [112], was used to normalize microarray intensity values across batches with default parameters (Fig. S3A). DOQTL was applied to the normalized microarray intensity values at each SNP probe to determine the founder strain that the SNP is most likely inherited from, i.e. the founder haplotype, as follows. Two founder haplotypes, referred to as founder diplotype, are possible at each SNP locus, giving a total of 36 possible founder diplotypes for a population generated from 8 founder strains: 8 homozygous diplotypes and 28 heterozygous diplotypes. DOQTL uses the intensity of each SNP probe to generate a probabilistic estimate for each diplotype state at each SNP in each DO mouse using a hidden Markov model, where the hidden states are the diplotype states. DOQTL assigned each SNP locus to the diplotype state with the highest posterior probability.

#### Individual mouse genome reconstruction

Seqnature [68] was used to construct an individual diploid genome for each DO mouse using founder haplotypes that were inferred at the previous step. Recombination boundaries were defined as the genomic midpoint between neighboring SNPs that were assigned to different founder diplotypes. For each recombination block, Seqnature was used to recapitulate the founder genome of interest by incorporating high quality SNPs and small indels (<100 bases) that were found in that particular genome, downloaded as a vcf file from the Sanger Mouse Genome project (release 1211, [62]).

#### RNA-seq read mapping

Tophat2 [113, 114] was used with default parameters to map RNA-seq reads from each individual DO mouse liver sample to that mouse’s diploid genome, which was comprised of both paternal and maternal allele sequences (Fig. S3A).

### Other mouse liver RNA-seq samples and data analyses

Three publicly available liver gene expression datasets were downloaded for the eight DO mouse founder strains: 1) a microarray dataset comprised of 96 samples, six replicates per sex for each strain, assayed on the Illumina Sentrix Mouse-6 V1.1 platform, at http://cgd.jax.org/gem/strainsurvey26/v1; 2) an RNA-seq dataset comprised of 128 male liver samples, six livers per strain from each of two diets, standard chow and high fat diet (GEO accession GSE45684) [67, 68]; and 3) an RNA-seq dataset comprised of 12 male and 12 female mouse livers for strain C57Bl/6J (GEO accession GSE59222) [3]. Tophat2 was used with default parameters to map each RNA-seq sample to its respective strain’s genome, generated by Seqnature based on SNPs/indels retrieved from the Sanger Mouse Genome project (release 1211) [62]. RNA-seq data for livers from the CD-1 (ICR) mouse strain, consisting of three pools of male and three pools female liver samples, was downloaded from GEO accession GSE98586 [34] and then mapped to the reference mouse genome using Tophat2.

### Expression quantification

Gene expression was quantified using featureCounts [115] to count sequence reads that overlap any exon by at least one bp. For RNA-seq datasets that were mapped to a haploid genome, e.g. the C57Bl/6J reference mouse genome and the eight DO mouse founder strains, only uniquely mapping reads were used. For DO mouse datasets, which were mapped to the reconstructed diploid genome of each individual mouse, the best mapped location for each read was used. The restriction of using only unique mapped reads was not imposed, as it would exclude reads that map to genomic regions where the paternal and maternal alleles are the same, which in many cases would substantially underestimate expression of a gene. Gene expression levels for DO mouse RNA-seq data were based on the total number of reads that overlap the counted regions in either the paternal or maternal allele. Read counts were then transformed to fragments per kilobase of exon per million reads mapped (FPKM) for downstream analysis. For FPKM calculation on diploid genomes, exon lengths were based on the average length of exons for the two alleles of each gene.

### Sex-specificity of gene expression

Three liver expression datasets, described above, were used to establish the sex-specificity of protein-coding genes: 1) male and female CD1 mouse liver RNA-seq dataset; 2) male and female C57Bl/6J mouse liver RNA-seq dataset; and 3) male and female mouse liver Illumina Sentrix Mouse-6 V1.1 expression microarray dataset, encompassing each of the eight DO mouse founder strains. edgeR [116] was used to perform differential analysis for the two RNA-seq datasets; and limma [117] with default parameters was used for the microarray dataset. For genes with multiple microarray probes, we chose the probe with the lowest adjusted p-value. We considered a total of 1,196 sex-specific genes, as follows. A total of 1,033 protein-coding genes were designated sex-specific (Table S1) based on the following criteria: male/female gene expression |fold-change| > 2 at FDR < 0.05 in at least one RNA-seq dataset, or |fold-change| > 1.5 at FDR < 0.05 in at least one of the microarray datasets. A set of 168 multi-exonic, intergenic liver-expressed lncRNAs designated sex-specific based on their significant sex-biased expression in one or more of the 8 RNA-seq datasets examined previously (male/female |fold-change| > 4 at FDR < 0.05; see listing in Supplemental Table 2A, column AB, of [20]), of which 163 were expressed in at least one DO liver sample. For each gene, the sex specificity was based on its sex bias in the strain with the largest |fold-change| for male/female expression ratio. Where indicated, our analyses were limited to 471 sex-specific protein-coding genes for which microarray expression data is available for all eight DO founder strains in both sexes.

### Variability of gene expression across DO founder strains

RNA-seq data for n = 8 male livers in each DO founder mouse strain was used to determine the variability across, and within, each founder strains, as follows: Inter-strain variability (Gene X) = [standard deviation of (mean expression value of Gene X in each of the 8 strains)] / [average of (set of mean expression values for Gene X in each of the 8 strains)]. Intra-strain variability (Gene X) = [standard deviation of Gene X expression across n = 8 male livers in strain *s*] / [mean of Gene X expression in n = 8 male livers in strain *s*].

### eQTL analysis

eQTLs were identified using liver gene expression data 219 male and 219 female DO mouse livers (Fig. S3A, Table S2). We used the additive haplotype model in DOQTL [112] to perform QTL mapping by regressing each gene’s expression level based on the estimates of each founder strain’s contribution, i.e., the founder diplotype probability, at each SNP marker. Adjustments to account for relatedness amongst DO mice, as well as batch, sex, diet, and interaction between sex and diet were included in the regression analysis as additive covariates, written as the following in R: model.matrix(~sex + diet + diet*sex + generation + batch, data = data). The strength of the association between gene expression and genotype is given as a likelihood ratio (LOD score, logarithm of the odds score), which is the −log10(p-value) when comparing the full model to the null model, where the null model excludes diplotype probabilities. QTL mapping also gives eight regression coefficients, whose magnitudes reflect the effect of the founder alleles at each SNP marker. A positive regression coefficient indicates the genetic variant is associated with high expression of the gene of interest, and a negative regression coefficient indicates the genetic variant is associated with low expression of the gene of interest. For each eQTL, the regulating strain (i.e., the strain with the largest alteration in gene expression due to genetic factors) was defined as the DO founder strain whose regression coefficient had the largest absolute value. A Bayesian credible interval, defined as 95% of the region under the LOD^10 curve, as implemented in DOQTL [112], was defined for each eQTL.Tshis interval delineates the genomic location where the highest association occurs, as defined by the uppermost 5% area under the peak of the LOD score curve.

#### Significance level

A genome-wide p-value for each association was determined by carrying out 1,000 permutations on the gene expression data. To assemble a list of associations between gene expression and SNPs, the SNP with the highest LOD score for each gene was recorded; subsequent high-scoring SNPs were retained if their p-value was < 0.05 and if they were located on a different chromosome than the SNP with the highest LOD score. An FDR correction was then applied to the genome-wide p-values. Any eQTL with FDR < 0.05 was deemed significant. To identify genetic variants whose association with gene expression was only found in one sex, or was stronger in one sex than in the other, we repeated the eQTL mapping analysis twice more, once using only the 219 male DO liver samples, and a second time using only the 219 female DO liver samples.

#### Gene expression pre-processing

Gene expression levels, in units of FPKM, were transformed into normal scores using the inverse normal transformation in DOQTL [112] prior to their use for eQTL mapping. eQTL mapping was carried out for protein-coding genes and for intergenic, multi-exonic lncRNA genes that were expressed in at least one liver sample, namely: 20,559 genes (18,543 protein-coding genes and multi-exonic 2,016 lncRNA genes) when analyzing the 438 male and female DO livers together as a single set; 20,268 genes (18,275 protein-coding and 1,993 lncRNA genes) when analyzing only the 219 male DO livers; and 20,190 genes (18,207 protein-coding and 1,983 lncRNA genes) when analyzing only the 219 female DO livers.

#### Interpolating missing genotype information

To preserve the extra information that the higher density SNP array provides, QTL mapping was performed on 64,713 SNPs markers (Table S2A), which corresponds to the union of SNP markers from the two genotyping arrays used after removal of SNP probes with no differentiating information across the eight strains [111]. For missing SNP probes, i.e. SNPs that were unique to one type of array, diplotype probabilities were assigned from the nearest measured SNP. All of the SNP probes that were analyzed are located on autosomes or on chrX.

#### Novelty of eQTLs

An eQTL was considered novel if its 95% Bayesian credible interval did not overlap by at least one bp the associated variant region of a published eQTL for the same gene name or RefSeq accession number.

#### *Cis* and *trans* eQTLs

An eQTL was designated *cis* if the eQTL’s 95% Bayesian credible interval overlapped (by at least 1 bp) the TAD region that contains the transcription start site of the gene regulated by the eQTL; otherwise it was designated a *trans* eQTL. TAD coordinates were those listed for mouse liver in [71].

### Categorization of eQTLs with a sex-bias in genetic association

Sex-biased genetic associations were defined for 360 autosomal eQTLs that are significant in either male liver or female liver samples (see Table S4A, columns D, F, N and O), based on LOD score calculated using male only DO liver samples *minus* LOD score calculated using female only DO liver samples. Each such sex-dependent eQTL was assigned to one of eight categories based on three criteria: the sex-specificity of the gene it is associated with, the sex-bias of its genetic association (LOD score in males minus LOD score in females), and its regression coefficient in the regulating strain and in the sex where the eQTL LOD score is higher (Table S5). A decrease (or an increase) in sex specificity was inferred from a negative (or a positive) regression coefficient, which indicates that the gene is predicted to be expressed at a lower (or higher) level in the regulating strain as compared to its expression in the other DO founder strains based on the regression model. For example: category #1 is comprised of eQTLs associated with male-specific genes that show stronger regulation in male liver (LOD score in males > LOD score in females) and a negative regression coefficient in the regulating strain, indicating a decrease in sex specificity compared to what it would have been without the eQTL; while category #8 is comprised of eQTLs associated with female-specific genes that show stronger regulation in female liver (LOD score in females > LOD score in males) and a positive regression coefficient in the regulating strain, indicating an increase in sex specificity compared to what it would have been without the eQTL (Table S5).

### Transcription factor binding sites and DHS (open chromatin regions) with strain-specific SNPs/indels at eQTLs

High quality SNPs/indels in the eight DO mouse founder strains were downloaded from the Sanger Mouse Genome project (release 1211) [62]. SNPs/indels that occur in only one founder strain were designated strain-specific variants, and were extracted from a variant call format (vcf) file downloaded from https://www.sanger.ac.uk/sanger/Mouse_SnpViewer/rel-1211 (release 1211) using a custom R-script provided in File S1. We selected for analysis DHS and transcription factor binding site ChIP-seq peaks that are within each eQTL region (95% Bayesian Credible interval, based on the liver sample set that gives the highest LOD score), and whose peak region contains a strain-specific SNP/indel for the matching regulating strain. For these analyses, we considered eQTL regions that are < 3 MB in width (8,254 of all 12,886 autosomal eQTLs). Further, eQTLs identified for genes with |male/female| expression > 4 were designated an eQTL for a sex-specific gene, and eQTLs identified for genes with |male/female| < 1.2 were designated an eQTL for a sex-independent gene, based on male/female expression ratios determined by edgeR. The number of eQTLs included in the analysis of DHS and transcription factor binding sites ranged from 17 and 2,793, and is specified for each analysis in the x-axis labels of Fig. 4. For each eQTL region, we counted the fraction of DHS or ChIP-seq binding sites for each transcription factor analyzed that are male-specific, female-specific or sex-independent, as defined in the published studies for STAT5 [32] and BCL6 (Table S11 of [21]), and for the set of ~72,000 liver DHS from [33], of which 4,644 DHS were designated male-biased DHS and 2,814 were designated female-biased DHS based on reanalysis of the published raw data performed by Gracia Bonila of this laboratory. For each eQTL considered, we computed the fraction of all male-specific, female-specific and sex-independent DHS, or transcription factor binding sites, within the eQTL region that contain a SNP/indel specific to the regulated strain for that eQTL (Fig. S6), and then compared the distributions of those fractions to those of the other sets of eQTLs being considered, as shown along the x-axis of Fig. 4. Significance was determined by Wilcoxon test.

### Co-regulated gene clusters

Each eQTL region was centered at the SNP marker with the highest LOD score, i.e. the eQTL peak. eQTL region boundaries were set at the second SNP marker from the center; thus, each eQTL region contains five SNP markers. eQTL regions with the same regulating strain whose boundaries overlap by at least one bp were assigned to the same co-regulated gene cluster. The genomic region encompassing overlapping eQTLs was computed two ways: 1) based on the intersection of the regions defined by the five SNP markers of each eQTL; and 2) based on the union of the 95% percent Bayesian intervals of each of the overlapping eQTLs. Of note, the 95% percent Bayesian interval may be smaller or larger than the five SNP region, and the union of 95% Bayesian intervals may also be smaller than the intersection of the five SNP regions, depending on how large and overlapping the 95% Bayesian intervals are. Co-regulated gene clusters were identified based on eQTLs discovered in all DO liver samples, and separately, male only DO liver samples, and female only DO liver samples. This analysis considered eQTLs for genes that are expressed (FPKM > 0) in at least 25% of each set of liver samples, i.e., all DO livers, male only DO livers, or female only DO livers.

### Other analyses

The var function in R was used to quantify liver expression variation for sex-specific genes across livers from 112 chow diet-fed male DO mice, and separately across livers of 110 chow diet-fed female DO mice. Sex-specific genes belonging to each of four classes, defined by their responses to hypophysectomy, were from Table S3 of [30]. The ComBat function in the sva R package [118] was used with default parameters to remove from the gene expression data (in units of log2(FPKM+1)) a batch effect that correlated with the liver sample’s GEO accession number. Pearson correlation was then used to calculate pairwise gene expression correlations. PCA was implemented using the prcomp function in R using log2, centered normalized microarray intensity values. DAVID functional annotation (https://david.ncifcrf.gov/) [64, 65] was used to discover enriched KEGG pathways by inputting the official gene symbols for the sex-specific protein-coding genes discovered in each strain. Pathways with adjusted P (Benjamini) < 0.05 were deemed significant. liftOver [119] was used to convert genomic regions into mouse mm9 coordinates using default parameters, and bioDBnet:db2db [120] was used to convert Ensembl gene identifiers to RefSeq accession numbers.

## Supporting information

**Fig. S1. High variability of the sex-specificity of gene expression in individual livers of CD-1 mice and DO mouse founder strains for two examples of sex-specific genes.** Gene expression in individual mouse livers across strains: male and female CD-1 mice (first row; *left*), male and female C57BL/6J mice (first row, *right*), male and female DO mice fed a standard chow diet (*second row*), or fed a high fat diet (*third row*). The fourth row shows box plots of gene expression level (FPKM) based on 128 individual male livers for DO founder mice fed a standard chow diet (*left*), or fed a high fat diet (*right*), and for male and female DO founder strain mice (fifth row; *left*). The first four rows present gene expression values determined by RNA-seq (FPKM values), while the gene expression on the fifth row was determined by microarray analysis. Male/Female expression ratios across the DO founder strains based on the microarray dataset are also presented (fifth row; *right*). Examples shown are for *Cyp4a12b*, a male-specific gene (A), where male-biased expression is reduced or lost in PWK/PhJ mice, and for *Cyp2c39,* a female-specific gene (B), where female-biased expression is lost in C57BL/6J mice.

**Fig. S2. Numbers of sex-specific protein-coding genes in each DO founder strain.** Shown are the number of sex-specific protein-coding genes (**A**) or number of highly sex-specific protein-coding genes **(B)** (male/female |fold-change| > 4 at FDR < 0.05) in at least one DO mouse founder strain based on the microarray dataset) for each founder mouse strain.

**Fig. S3. Analysis of SNP array and RNA-seq data for each DO mouse liver sample: Schematic overview (A), and properties of autosomal eQTLs in DO mouse liver (B).** Shown in (B) are the percentages of autosomal eQTLs that are associated with increased expression (white bars) or decreased expression (colored bars) in the respective regulating founder mouse strain. These data are based on 625, 672, 574, 593, 616, 3,085, 3,074 and 1,086 autosomal eQTLs whose regulating strain is A/J, C57BL/6J, 129S1/SvlmJ, NOD/ShiLtJ, NZO/HILtJ, CAST/EiJ, PWK/PhJ, or WSB/EiJ, respectively. Data are based on the 10,325 autosomal eQTLs (Table 1) discovered when the set of all DO liver samples were used for eQTL discovery.

**Fig. S4. Examples of sex-specific genes with stronger genetic association in male or female liver.** For each gene, the figure shows the gene expression patterns across strains (see Fig. S1 for details), and genome-wide eQTL scans using the three different DO liver datasets, where the horizontal red line marks the P < 0.05 significance cutoff for eQTL significance based on the permutation test. Also shown are regression coefficients across the chromosome that contains the significant eQTL peak, as follows: Regression coefficients (top of each panel) and LOD scores (log10(p-value); bottom of each panel) across the chromosome that has a significant eQTL peak in male (left) or female (right) mouse liver, as marked at bottom. Shaded area in the LOD score plot indicates the 95% Bayesian credible interval for each eQTL. **(A)** *Cav1* shows strong male-specific expression in the PWK/PhJ founder strain and in a subset of male DO mouse livers. Its strong eQTL is seen in male livers only and is associated with the PWK/PhJ strain. **(B)** *Hsd3b5* shows male-biased expression in CD-1 mice and multiple DO founder strains; its expression is repressed in female C57BL/6J mice.

**Fig. S5. Examples of sex-specific genes in categories # 1-8, described in Fig. 3.** For each gene, the figure shows gene expression patterns across strains, genome-wide eQTL scan, and regression coefficients at the chromosome with a significant peak, as described in the legend to Fig. S1.

**Fig. S6. Enrichment of sex-specific regulatory elements with strain-specific SNPs/indels at eQTLs for sex-specific genes.** In the example shown, 5 of the 8 regulatory elements (i.e., ChIP-seq binding sites) within the eQTL region shown are male-specific binding sites, three of which contain either SNPs or Indels specific for the regulating strain. A fourth regulatory element with a strain-specific SNP is a sex-independent binding site. One strain-specific SNP is not in a regulatory element, and so is excluded from the analyses shown in Fig. 4, as are the four reguatory elements without any strain-specific SNPs/indels.

**Fig. S7. Regression coefficients (top figure in each panel) and LOD scores (bottom figure in each panel) for genes shown in Fig. 5A.**

**Fig. S8. Genome-wide LOD score plots for genes that are located in different chromosomes in the co-regulated cluster depicted in Fig. 5A**.

**Fig. S9. Genome-wide LOD score plots the *Sult2a1*, *Sult2a3* and *Sult2a5* genes**.

**Fig. S10. Strain-specific SNPs/indels proposed to contribute to the loss of sex-specific gene expression for *Cyp2b9* in female 129S1/SvlmJ liver**. (A) distribution of gene expression level of *Cyp2b9* across individual DO mouse livers (Table S2) was used to discover eQTLs stratified by the genotype assigned at the the SNP marker with the highest LOD score. (B) A female-biased STAT5 binding sites (fourth red arrow from the top) at a female-biased DHS (third red arrow from the top), containing two 129S1/SvlmJ-specific SNPs/indels (two top arrows), located within the eQTL region for *Cyp2b9*.

**File S1.** Files for custom R-script used to extract strain-specific SNPs/indels from variant call format file downloaded from https://www.sanger.ac.uk/sanger/Mouse_SnpViewer/rel-1211 (release 1211).

## Acknowledgments

We thank Gracia Bonilla of this laboratory for providing lists of male-biased and female-biased DHS used in Fig. 4, and Dr. Daniel Gatti of The Jackson Laboratory for providing access to the DO mouse SNP array and liver expression datasets used in this study.

